# T1R2/T1R3 Identified as the Aβ Receptor: A New Mechanism Linking Aβ42/40 Ratio to Neuronal Hyperexcitability in Alzheimer’s Disease

**DOI:** 10.64898/2026.09.25.754367

**Authors:** Mitsuyoshi L. Saito, Mariko R. Saito

**Author notes:** Address: Mitsuyoshi L. Saito, Ion Chat Research Corp., Tokorozawa city, Saitama, 359-1146, Japan.

## Abstract

Early neuronal hyperexcitability is a hallmark of Alzheimer’s disease (AD) and correlates with cognitive decline. A key diagnostic indicator is the reduction in soluble Aβ42/40 ratio - from ∼10% in cognitively normal individuals to ∼5% in AD - which reliably predicts progression to dementia. Although widely used as a biomarker, whether this ratio plays an active signaling role has remained unclear. Building on our prior finding that T1R3 is required for Aβ25–35–mediated enhancement of voltage-gated sodium current (INa), we now show that the AD-associated 5% Aβ42/40 ratio functions as an active ligand that increases INa, whereas the physiological ∼10% ratio is inert. This effect requires T1R2 and T1R3, but not T1R1, identifying T1R2/T1R3 as the receptor responsible for Aβ ratio sensing. Human cell lines expressing Nav1.1, Nav1.5, or Nav1.6, as well as differentiated SH-SY5Y cells, all respond selectively to the active ratio. Rodent NG108-15 cells are natively insensitive, but expression of human T1R3 confers responsiveness, indicating a human-specific detection mechanism. These findings identify T1R2/T1R3 as the receptor that detects the Aβ42/40 ratio, link early Aβ changes to hyperexcitability, and point to a therapeutic target for restoring early neuronal function. The human-specific nature of this pathway motivates development of humanized T1R2/T1R3 knock-in mice for in vivo validation.

**Significance Statement:** In Alzheimer’s disease progression, neuronal hyperexcitability is one of the underlying causes, yet its molecular mechanism remains unknown. Here we report that the pathological Aβ42/40 ratio, a cerebrospinal fluid biomarker of Alzheimer’s disease, enhances voltage-gated sodium current, the major driving force of action potential generation. We identify the receptor responsible for this effect as an extra-oral sweet taste receptor, T1R2/T1R3. This mechanism is human-specific because rodent cells are insensitive to the pathological Aβ42/40 ratio, providing a molecular clue to why rodent models fail to recapitulate early Alzheimer’s disease. These findings identify T1R2/T1R3 as a molecular entry point at which the earliest biochemical change in AD is converted into neuronal dysfunction, making it a valid target for therapeutic intervention.

## Introduction

Alzheimer’s disease (AD) begins with subtle but widespread alterations in neuronal excitability that arise years before overt neurodegeneration or cognitive decline. Hyperexcitability is detectable in patients with mild cognitive impairment (MCI), and this hippocampal hyperactivity appears to precede overt amyloid deposition (1). Among the earliest biochemical abnormalities is a reduction in the cerebrospinal fluid (CSF) Aβ42/40 ratio, which precedes amyloid deposition, shifts from the physiological ∼10% ratio to the ∼5% ratio characteristic of AD, and is already reduced at the MCI stage (2, 3). Although widely used as a biomarker, whether this ratio plays an active signaling role has remained unknown. This uncertainty is underscored by the paradox that endogenous Aβ at picomolar concentrations is required for normal synaptic plasticity and long-term potentiation (LTP) (4), suggesting that Aβ participates in a regulated receptor-mediated pathway whose disruption may contribute to early AD-associated hyperexcitability.

A major unresolved question is the identity of the receptor or molecular sensor that detects physiological Aβ signals. The Aβ42/40 ratio is tightly regulated in healthy individuals (5, 6, 7), and small deviations from this ratio correlate with early cognitive changes (8). Yet no receptor has been shown to selectively respond to this ratio, and no mechanism has been established to explain how ratio sensing could influence neuronal excitability. Compounding this gap, although several rodent AD models exhibit hyperexcitability at later stages of amyloid pathology, they do not recapitulate the early hyperexcitability observed in human MCI, suggesting that a human-specific receptor mechanism may be required to link early Aβ changes to neuronal dysfunction (8).

We previously demonstrated that INa in HEK and SH-SY5Y cells is enhanced by Aβ25–35 and L-glutamine through a T1R3-dependent mechanism, in which these cells endogenously express T1R1, T1R2, and T1R3 (9). Based on these observations, we asked whether the T1R2/T1R3 heterodimer responds to mixtures of full-length Aβ40 and Aβ42 at the physiological ∼10% and AD-associated ∼5% Aβ42/40 ratios, and whether it can distinguish between these two ratios in their ability to enhance INa. The T1R2/T1R3 complex - the canonical sweet taste receptor (10, 11, 12) - is broadly expressed in the brain, including neurons and astrocytes (13). Using patch-clamp recordings in sodium-channel-expressing human cell lines and differentiated SH-SY5Y cells, we found that the 10% Aβ42/40 ratio behaves as an inactive ligand, whereas the 5% ratio acts as an active ligand that increases INa amplitude. Gene-specific silencing experiments revealed that loss of either T1R2 or T1R3 abolishes the 5% ratio-induced INa enhancement, whereas T1R1 silencing leaves the response intact. These results identify T1R2/T1R3 as the molecular sensor that discriminates Aβ42/40 ratios.

We further show that the rodent NG108-15 cell line (14) is natively insensitive to the active 5% ligand, although it responds to L-glutamine with enhanced INa as seen in human cells (9). However, expression of human T1R3 confers responsiveness to the 5% ratio, demonstrating a human-specific mechanism. Our results demonstrate that the AD-associated shift in the Aβ42/40 ratio is not merely a diagnostic biomarker but a functional ligand that enhances INa. Identifying T1R2/T1R3 as the receptor for this ligand provides a mechanistic basis for early hyperexcitability and highlights a new therapeutic target. This species-restricted mechanism also creates the opportunity to generate humanized T1R2/T1R3 knock-in mice, enabling in vivo studies of early excitability changes.

## Results

### Aβ42/40 at the 5% ratio enhances sodium current amplitude in human SH-SY5Y neurons

To determine whether full-length amyloid-β (Aβ) peptides modulate voltage-gated sodium channels (Nav), we performed whole-cell voltage-clamp recordings in differentiated human SH-SY5Y neuroblastoma cells. Aβ₁₋₄₂/Aβ₁₋₄₀ mixtures were prepared at concentrations matching diagnostically measured cerebrospinal fluid levels. The Alzheimer’s disease–associated 5% ratio (ADC) contained 123.8 pM Aβ₁₋₄₂ and 2.5 nM Aβ₁₋₄₀, whereas the physiological 10% ratio (NDC) contained 254.5 pM Aβ₁₋₄₂ and 2.3 nM Aβ₁₋₄₀ (3). SH-SY5Y cells endogenously express T1R1, T1R2, and T1R3 (9), consistent with a potential role for the T1R2/T1R3 receptor in Aβ-dependent modulation of excitability.

Application of the 5% Aβ42/40 ratio significantly increased peak inward sodium current (INa) to 128.8 ± 4.5% of control (n = 8, p = 0.0004; Fig. 1a1, a2), with enhancement observed in every cell. In contrast, the 10% ratio produced no significant change (n = 5, p > 0.05; Fig. 1b1, b2). Thus, the 5% ratio acts as an active ligand capable of enhancing INa, whereas the 10% ratio is functionally inert under these conditions.

**Figure 1.**
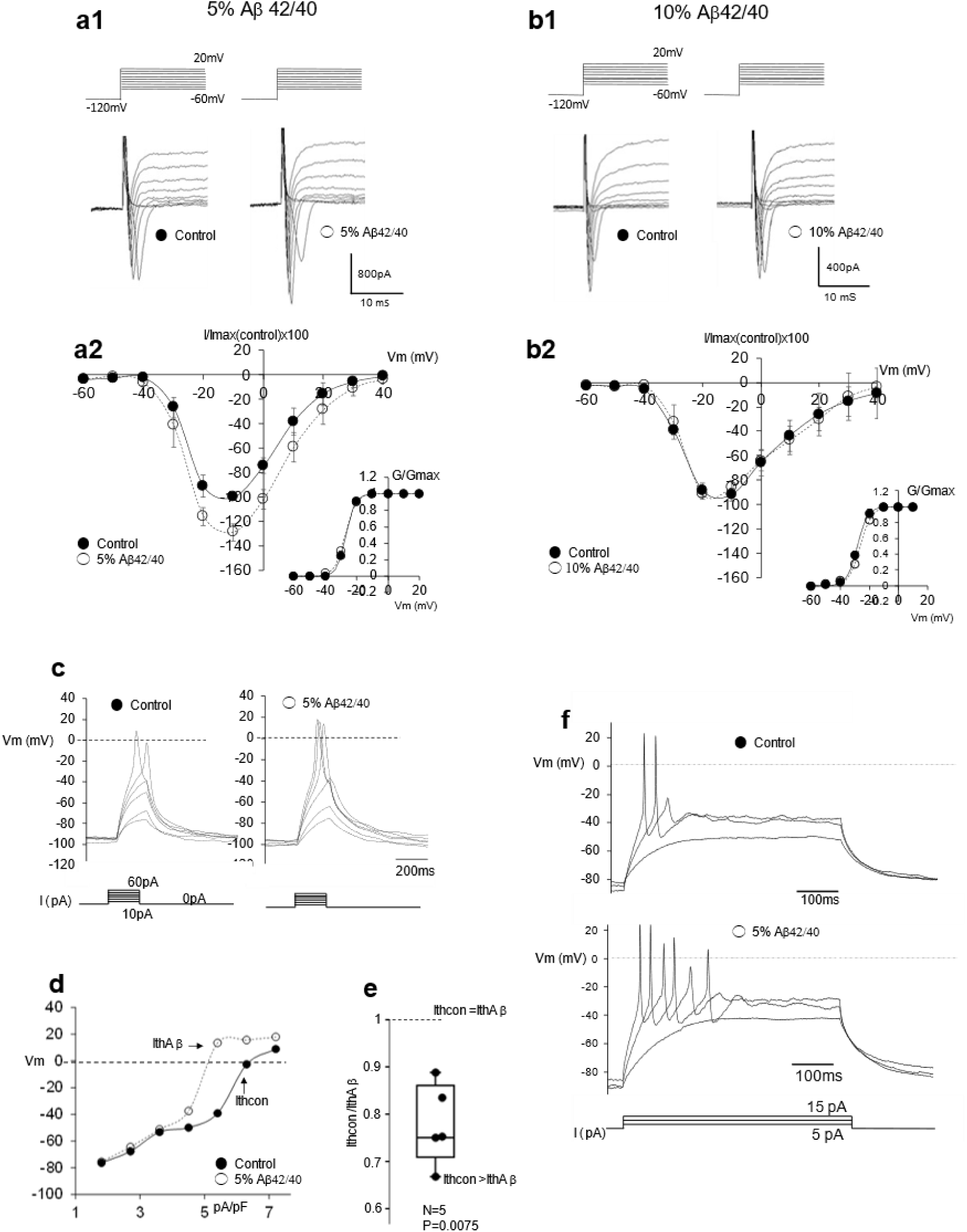
Aβ42/40 ratio-dependent modulation of sodium currents and excitability in differentiated SH-SY5Y neurons. Voltage clamp recordings. (a1) Representative whole-cell sodium currents evoked by voltage steps under control conditions (left) and after application of the 5% Aβ42/40 ratio (right). (a2) Averaged I-V relationships (mean ± SEM) showing a significant increase in peak INa after 5% Aβ42/40 (n = 8). Inset: G-V curves showing no detectable shift in activation. (b1) Representative sodium currents recorded before (left) and after exposure to the 10% Aβ42/40 ratio (right). (b2) Averaged I-V relationships for the 10% ratio showing no enhancement of INa (n = 5). Current clamp recordings. (c) Action potentials evoked by incremental depolarizing current injections under control conditions (left) and after 5% Aβ42/40 exposure (right), showing reduced action potential threshold. (d) Voltage-current relationship from a representative cell demonstrating a leftward shift in firing threshold from control (filled circles) to after 5% Aβ42/40 (open circles). Arrows indicate Ith(control) and Ith(Aβ). (e) Across cells (n = 5), the ratio Ith(Aβ)/Ith(control) was <1 in every case, indicating a consistent reduction in current required to reach spike threshold. (f) Representative repetitive firing traces showing increased firing frequency after 5% Aβ42/40.

G–V relationships (Fig. 1a2, b2) showed no detectable shifts in voltage dependence, confirmed by unchanged half-activation voltages (Vh) (Fig. 2g2). The 5% ratio therefore increases INa amplitude without altering channel gating.

**Figure 2.**
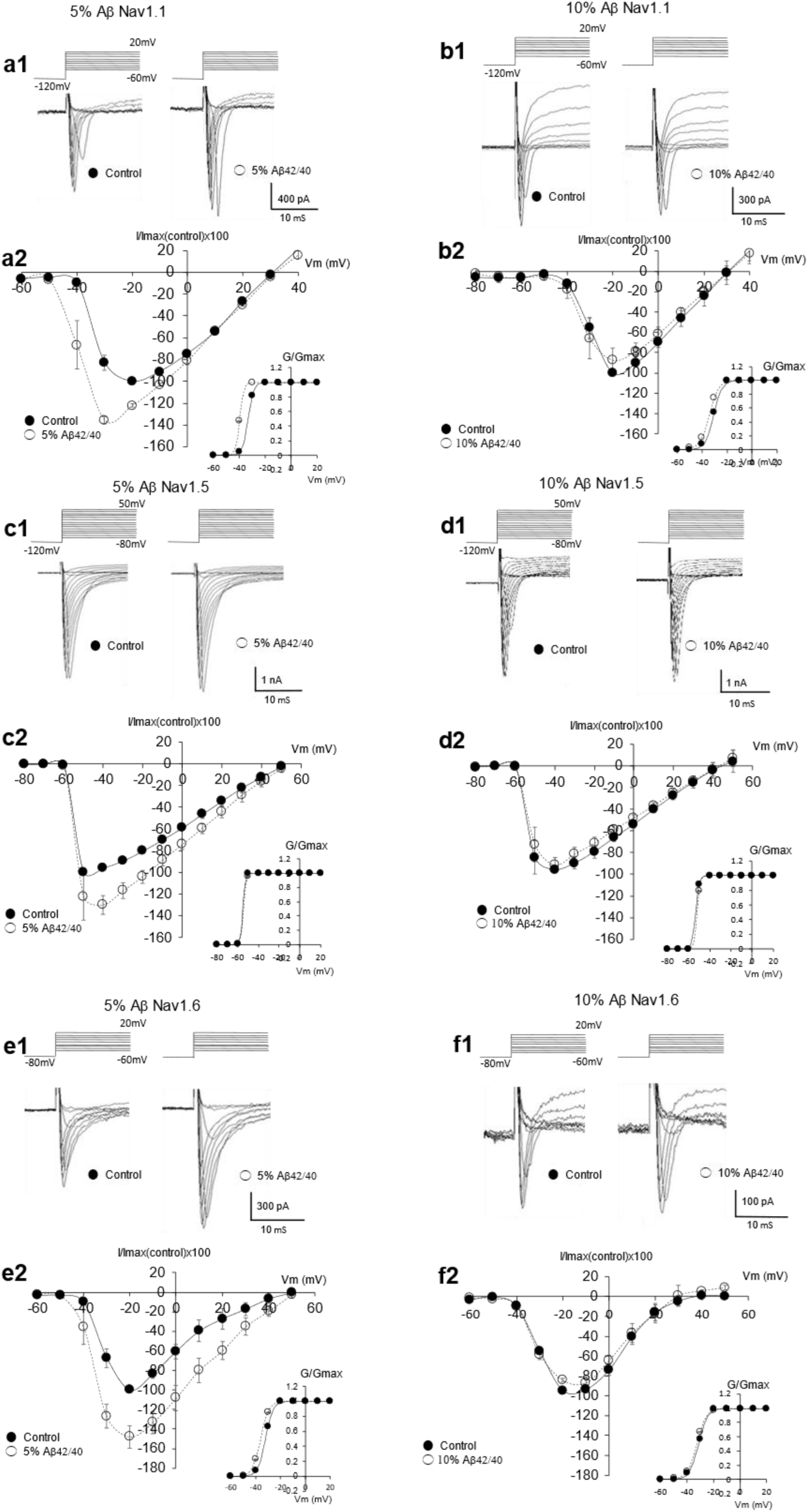
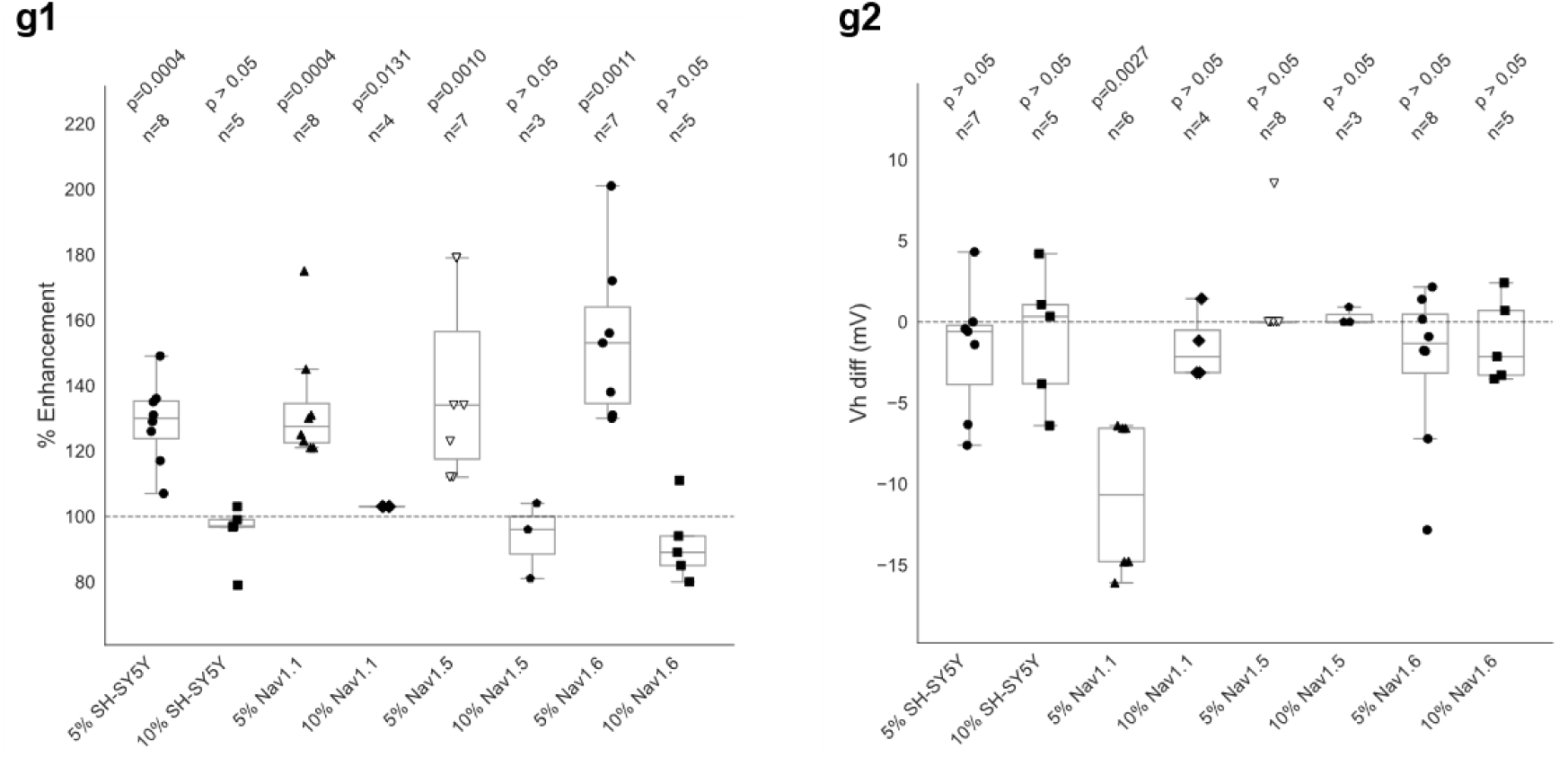
The 5% Aβ42/40 ratio enhances INa across Nav1.1, Nav1.5, and Nav1.6 channels expressed in HEK293 or SH-SY5Y cells. Representative whole-cell sodium current traces recorded from cells expressing Nav1.1 (a1), Nav1.5 (c1), or Nav1.6 (e1) under control conditions (left trace in each pair) and after application of the 5% Aβ42/40 ratio (right trace). A second pair of traces is shown for each isoform comparing control and 10% Aβ42/40. Averaged I-V relationships (mean ± SEM) for Nav1.1 (a2), Nav1.5 (c2), and Nav1.6 (e2) under control (filled circles) and 5% Aβ42/40 (open circles). Insets: corresponding Boltzmann G-V fits. Representative traces for the 10% ratio are shown for Nav1.1 (b1), Nav1.5 (d1), and Nav1.6 (f1), with corresponding averaged I-V plots (b2, d2, f2) and G-V insets. Sample sizes: 5% ratio — Nav1.1 (n = 8), Nav1.5 (n = 7), Nav1.6 (n = 7); 10% ratio — Nav1.1 (n = 4), Nav1.5 (n = 3), Nav1.6 (n = 5). (g1) Box-and-whisker plots summarizing percent enhancement of peak INa for each Nav isoform, including SH-SY5Y (Fig. 1a2, b2), showing robust increases at 5% and minimal changes at 10%. (g2) Box-and-whisker plots of Vh differences (Vh diff) between peptide and control conditions. Only Nav1.1 exhibited a significant shift at the 5% ratio; Nav1.5 and Nav1.6 showed no consistent change. Because voltage steps were applied in 10 mV increments, slope factors were not determined.

### The 5% Aβ42/40 ratio increases Nav-dependent excitability in human SH-SY5Y neurons

To test whether INa enhancement translates into altered excitability, we performed current-clamp recordings with incremental depolarizing current injections. Under control conditions, SH-SY5Y neurons required relatively large current steps to trigger action potentials. After exposure to the 5% ratio, action potentials were elicited at markedly lower current injections (Fig. 1c). Across five cells, the current threshold (Ith) was consistently reduced, with Ith(Aβ)/Ith(control) < 1 in every case (Fig. 1e).

In repetitively firing cells, the 5% ratio also increased action-potential firing frequency (Fig. 1f). Thus, the reduced Aβ42/40 ratio enhances INa, lowers spike threshold, and increases firing output, demonstrating that the ratio shift has a biological function beyond its role as a clinical biomarker.

### Aβ42/40 at the 5% ratio enhances INa across multiple Nav isoforms

To determine whether the 5% ratio acts through a shared upstream mechanism or directly modulates Nav channels, we examined its effects in cells expressing Nav1.1, Nav1.5, or Nav1.6. A receptor-mediated mechanism would be expected to produce qualitatively similar enhancement across isoforms, whereas direct channel interaction would likely yield isoform-specific variability. Application of the 5% ratio significantly increased INa in all isoforms tested: Nav1.1 (133.9 ± 6.5%, n = 8, p = 0.00041; Fig. 2a1, a2), Nav1.5 (139.0 ± 10.9%, n = 7, p = 0.00101; Fig. 2c1, c2), and Nav1.6 (154.4 ± 9.6%, n = 7, p = 0.00106; Fig. 2e1, e2). Enhancement magnitudes were comparable (Fig. 2g1). Voltage dependence remained largely unchanged, with only a small shift in Nav1.1 (Fig. 2g2). The 10% ratio produced no measurable enhancement in any isoform.

The conserved increase in INa across structurally distinct Nav channels supports a shared upstream mechanism, consistent with activation of a G-protein–coupled receptor rather than direct channel modulation.

### T1R2 and T1R3 are required for Aβ42/40-induced INa enhancement

Because HEK293 and SH-SY5Y cells express T1R1, T1R2, and T1R3 (9), we tested which protomers are required for the INa-enhancing effect of the 5% ratio. T1R1, T1R2, or T1R3 were individually silenced in Nav1.5-expressing HEK293 cells. Loss of T1R1 or T1R2 was confirmed by immunocytochemistry (Fig. 3d1, d2), and the T1R3 knockdown line was generated previously (9).

**Figure 3.**
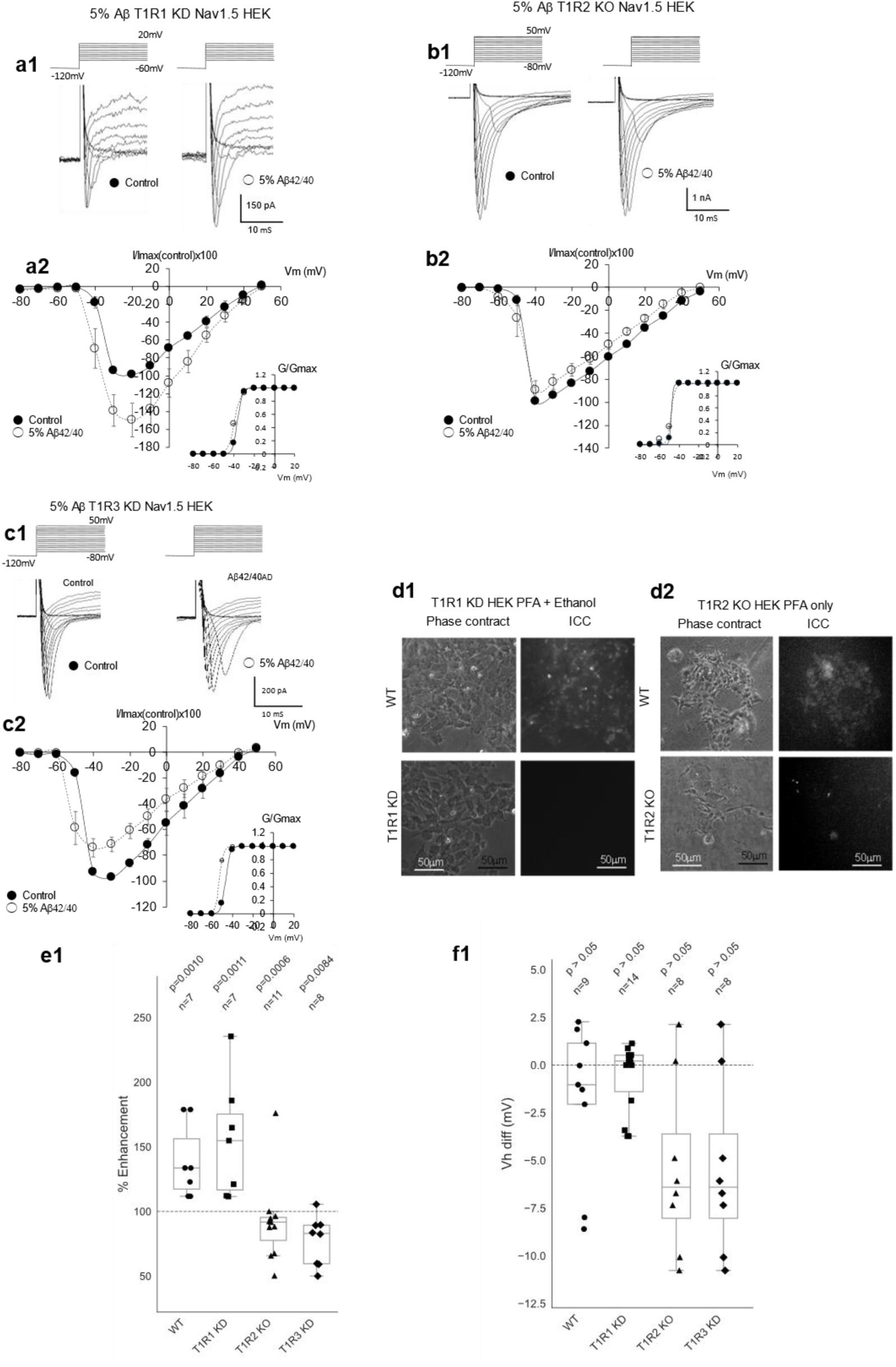
T1R2 and T1R3 are required for Aβ42/40-dependent enhancement of INa in Nav1.5-expressing HEK293 cells. (a1-c1) Representative INa traces and (a2-c2) normalized I-V relationships from Nav1.5-expressing HEK293 cells under control conditions (filled circles) and after application of the 5% Aβ42/40 ratio (open circles). Data are shown for T1R1 knockdown (KD) (a), T1R2 knockout (KO) (b), and T1R3 knockdown (KD) (c). Voltage clamp protocols match Fig. 2. Data are mean ± SEM. (d1-d2) Phase contrast (left) and immunocytochemistry (right) images confirming depletion of T1R1 (d1) or T1R2 (d2) in the respective KD/KO lines (upper panels: WT; lower panels: KD/KO). T1R3 KD validation was reported previously (9). (e1) Box-and-whisker plots summarizing peak INa enhancement (normalized to each experiment’s internal control): WT Nav1.5: 139.0 ± 10.9% (n = 7, p = 0.00101); T1R1 KD: 155.1 ± 17.8% (n = 7, p = 0.00106); T1R2 KO: 91.9 ± 9.6% (n = 11, p = 0.00059); T1R3 KD: 77.4 ± 6.7% (n = 8, p = 0.00838). T1R2 KO and T1R3 KD cells showed no enhancement and instead exhibited significantly reduced normalized INa, indicating loss of Aβ42/40 responsiveness. Individual data points are overlaid. (e2) Box-and-whisker plots of Vh diff showing no significant shifts across KD/KO conditions. All statistical comparisons used the Mann-Whitney U test.

In T1R1-KD cells, the 5% ratio significantly enhanced INa (155.1 ± 17.2%, n = 7, p = 0.00106; Fig. 3a), indicating that T1R1 is not required. In contrast, T1R2-KO and T1R3-KD cells showed no enhancement and instead exhibited significant reductions in INa (T1R2-KO: 91.8 ± 9.6%, n = 11, p = 0.00059; T1R3-KD: 77.4 ± 6.74%, n = 8, p = 0.008438; Fig. 3e1). Voltage dependence remained stable across conditions (Fig. 3e2).

These findings demonstrate that both T1R2 and T1R3 are required for Aβ42/40-induced INa enhancement, identifying T1R2/T1R3 as the necessary receptor complex.

### Human T1R3 confers Aβ42/40 responsiveness to NG108-15 cells

To test whether rodent cells respond to the 5% ratio, we performed voltage-clamp recordings in the mouse neuroblastoma × rat glioma hybrid line NG108-15, which expresses Nav1.7 (14). The 5% ratio produced no change in INa (94.9 ± 3.62%, n = 5, p = 0.1188; Fig. 4b1, b2), indicating that endogenous rodent T1R receptors cannot detect this ligand.

**Figure 4.**
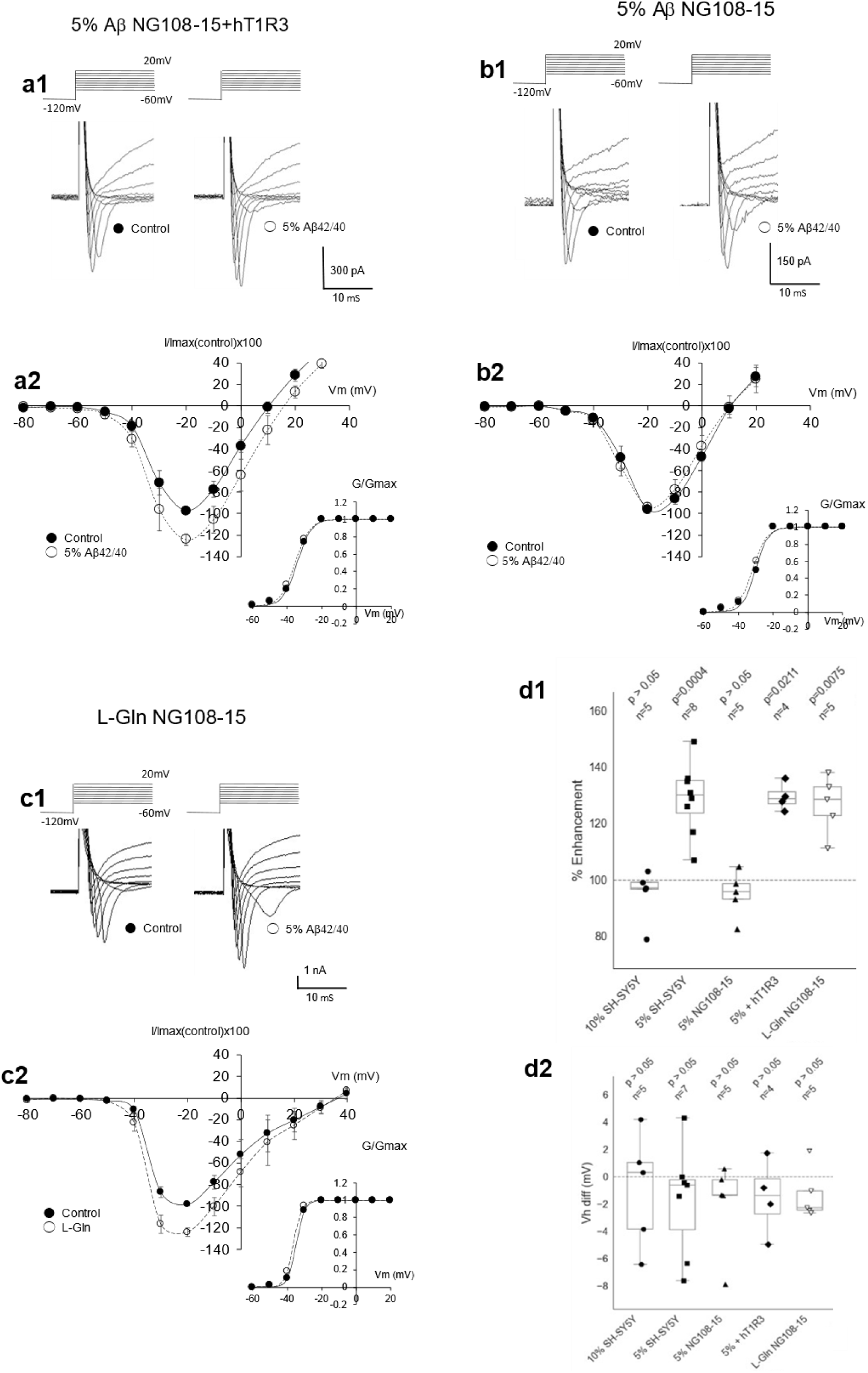
Human T1R3 confers Aβ42/40 responsiveness to undifferentiated NG108-15 cells. (a1) Representative INa traces from NG108-15 cells expressing human T1R3 under control conditions (left) and after application of the 5% Aβ42/40 ratio (right). (a2) Averaged I-V relationship (mean ± SEM) for NG108-15 + hT1R3 cells (n = 4). Inset: Boltzmann G-V fit. (b1) Representative INa traces from untransfected NG108-15 cells before (left) and after (right) exposure to the 5% ratio, showing no enhancement. (b2) Averaged I-V relationship (n = 5). Inset: G-V fit. (c1) Representative INa traces before (left) and after (right) exposure to 500 µM L-Glutamine (L-Gln). (c2) Averaged I-V relationship (n = 5). Inset: G-V fit. (d1) Box-and-whisker plots summarizing peak INa enhancement (normalized to internal control): SH-SY5Y: 128.7 ± 4.5% (n = 8, p = 0.00041) at 5%; 94.9 ± 4.1% (n = 5, p = 0.1188) at 10%; NG108-15: 94.9 ± 3.6% (n = 5, p = 0.1188) at 5%; NG108-15 + hT1R3: 129.5 ± 2.4% (n = 4, p = 0.02107) at 5%; L-Gln (500 µM): 126.7 ± 4.6% (n = 5, p = 0.00749). These results demonstrate that human T1R3 is sufficient to confer Aβ42/40 responsiveness in a rodent neuronal background and that rodent T1R2/T1R3 is functionally intact, as evidenced by L-Gln responsiveness. Individual data points are overlaid. (d2) Box-and-whisker plots of Vh diff showing no significant changes, indicating that differences in responsiveness reflect changes in current amplitude.

Expression of human T1R3 (hT1R3) conferred responsiveness: in hT1R3-transfected NG108-15 cells, the 5% ratio significantly enhanced INa (129.5 ± 2.4%, n = 4, p = 0.02107; Fig. 4a1, a2). This gain-of-function phenotype mirrors the response in human SH-SY5Y neurons and HEK293 cells expressing Nav1.1, Nav1.5, or Nav1.6.

L-glutamine enhanced INa in NG108-15 cells (126.7 ± 4.63%, n = 5, p = 0.00749; Fig. 4c1, c2), confirming that functional T1R2/T1R3 signaling is present. Thus, the lack of response to the 5% ratio reflects species-specific ligand recognition, not receptor absence.

Comparison across systems (Fig. 4d1) shows that in human SH-SY5Y cells, responsiveness is determined by ligand ratio (5% active, 10% inactive), whereas in NG108-15 cells, responsiveness is determined by receptor composition (rodent T1Rs inactive; hT1R3 active). Vh remained unchanged in all conditions (Fig. 4d2).

Together, these findings demonstrate that the 5% Aβ42/40 ratio is an intrinsically active ligand and that human T1R3 is the species-determining component for Aβ42/40 recognition.

## Discussion

### Part I. The Aβ42/40 Ratio as a Functional Signaling Ligand

Our findings redefine the conceptual role of the Aβ42/40 ratio. Rather than serving solely as a passive byproduct or biomarker of pathology, the ratio emerges as an active regulatory signal that directly modulates voltage-gated sodium channel (Nav) activity. We identify the extra-oral sweet taste receptor T1R2/T1R3 as the receptor complex that detects this signal, establishing a mechanistic link between pathological reductions in the Aβ42/40 ratio and early neuronal hyperexcitability. Importantly, this receptor-dependent mechanism is human-specific.

Rodent NG108-15 cells do not respond to the 5% Aβ42/40 ratio, indicating that the 5% ratio is a human-specific signal. Their endogenous T1R3 is functional, as shown by their sensitivity to saccharin (16), which activates mouse T1R2/T1R3 heterodimers or T1R3 homodimers (46), and by their significant INa enhancement in response to the T1R2/T1R3 agonist L-glutamine (L-Gln)(Fig. 4c1, c2), consistent with the effect observed in human cells (9). This shows that the intracellular machinery linking T1R2/T1R3 activation to INa enhancement is fully present in NG108-15 cells, even though the 5% Aβ42/40 ratio fails to activate it. Thus, the inability of rodent NG108-15 cells to detect the 5% Aβ42/40 ratio cannot be attributed to absent or non-functional T1R3. Instead, it reflects a species-specific inability of rodent T1R3 to recognize Aβ42/40 as a ligand. This conclusion is directly supported by our gain-of-function experiment: introducing human T1R3 into NG108-15 cells makes them responsive to the 5% ratio, producing INa enhancement comparable to human cells (Fig. 4a1, a2). This species specificity explains why rodent AD models fail to exhibit the early hyperexcitability observed in human MCI, and why the underlying signaling mechanism remained invisible.

This work builds on our previous finding that both L-Gln and Aβ25–35 enhance INa via T1R3 (9). In the present study, we demonstrate that full-length Aβ peptides, presented as mixed Aβ42/40 ratios, potentiate INa at clinically relevant picomolar concentrations (4). Given that T1R3 can heterodimerize with either T1R1 or T1R2, we next asked which pairing mediates Aβ ratio sensing. By individually silencing each protomer, we identified the T1R2/T1R3 complex as the required pairing for Aβ ratio sensing. Together, these findings indicate that this receptor complex functions as the primary sensor linking clinical shifts in the Aβ42/40 ratio (5–7) to early changes in neuronal excitability associated with the onset of Alzheimer’s disease.

### L-Glutamine and Aβ as a Dual-Ligand Sensory System

The observation that both L-Gln and the 5% Aβ42/40 ratio enhance INa through T1R2/T1R3 suggests that this receptor functions as a dual-ligand sensor. L-Gln is actively transported into synaptic vesicles by SLC6A17 (18), and its loss of function leads to significant cognitive deficits (17, 18), highlighting its importance in synaptic physiology. Given that L-Gln is a known ligand for this T1R–Nav pathway, bursts of synaptic activity likely trigger co-release of L-Gln along with L-glutamate, activating T1R2/T1R3 to enhance presynaptic INa. This response is accompanied by increased cellular capacitance (9), consistent with activity-dependent vesicle fusion that strengthens synaptic efficiency through a rapid, physiological feedback loop.

In contrast, Aβ is released into the interstitial space more slowly and persists longer than L-Gln (2). This allows Aβ-dependent enhancement of INa to provide a sustained modulatory effect that outlasts the initial bursting activity. Because rodents lack the specific 5% Aβ42/40 ratio-sensing layer found in humans, the resulting hyperexcitability is absent in rodent AD models unless severe Aβ aggregation is present (8). However, both species show significant learning impairments when SLC6A17 is compromised, indicating that L-Gln likely serves as the primary activator of the T1R2/T1R3 system, and that L-Gln-dependent activation represents an evolutionarily older, conserved component of the system.

The L-Gln - T1R3 pathway is therefore an evolutionarily conserved foundation. Humans, however, acquire an additional Aβ-dependent layer for long-lasting modulation, allowing continued T1R2/T1R3 activity after heightened synaptic activity subsides. This creates a more complex signaling landscape - one that is beneficial under physiological conditions but becomes a liability when the active 5% Aβ42/40 ratio remains chronically elevated.

### Part II. Resolving the “Biomarker Paradox” Through Conformational Invisibility

This model may help resolve a long-standing paradox: why Aβ42 appears to “fall” in the CSF of MCI patients before plaques are present. We propose that this decline reflects conformational invisibility. In early MCI, Aβ42 adopts compact oligomeric structures in which key epitopes are masked, rendering the peptide undetectable by standard immunoassays. This interpretation is supported by the observation that boiling SDS samples—denaturing and “unmasking” the protein—markedly increases Aβ42 immunoreactivity (21).

The formation of these oligomeric species may arise from the structural behavior of Aβ42 dimers, which display substantially greater conformational variability than Aβ40 homodimers (19, 20). The two hydrophobic residues unique to Aβ42 (Ile41 and Ala42) act as the mechanical trigger (22). Under physiological conditions (∼10% ratio), Aβ42 readily finds another Aβ42 to form homodimers. However, sustained neuronal activity elevates astrocytic Ca²⁺ (29), which is likely responsible for triggering Aβ40 release from perisynaptic astrocytic processes (25–27). This influx of monomeric Aβ40 could dilute local Aβ42, preventing Aβ42 from locating another Aβ42 partner. The system may behave analogously to super-cooled water: stable only until the environment shifts, at which point a single perturbation triggers an abrupt phase transition - Aβ42 snapping from its homodimeric state into the Aβ40:Aβ42 heterodimer, just as super-cooled water crystallizes into ice with a single tap.

In an Aβ40:Aβ42 heterodimer, the extra hydrophobic residues could impose curvature stress on the flexible 25 region backbone segment (23), creating a local “kink” that exposes residues 25–35 to solvent (24). This exposed region appears to correspond to the active domain previously shown to enhance INa through T1R3 (9), providing a structural basis for the functional equivalence between Aβ25–35 and the 5% Aβ42/40 ratio.

Our electrophysiological assay, using T1R2/T1R3, effectively functions as a biosensor for this hidden population. The fraction of Aβ42 that becomes invisible to diagnostic assays likely corresponds to the functionally active Aβ40:Aβ42 heterodimers driving neuronal hyperexcitability. Quantifying this invisible fraction may offer new insight into the etiology of Alzheimer’s disease.

### T1R2/T1R3 Ligand Handover: From Synaptic L-Gln to Active Aβ42/40

T1R2/T1R3 may integrate neuronal activity across distinct timescales through sequential engagement of two ligands. During LTP induction, synaptic vesicles release L-Gln (17, 18), and neuronal endosomes release Aβ42/40 at the physiological ∼10% ratio (28). L-Gln provides the initial fast signal, activating T1R2/T1R3 to promote microtubule stabilization and vesicular trafficking that enhance INa (9).

As discussed in Part II, sustained activity produces neurotransmitter spillover that elevates astrocytic Ca²⁺ (29) to the threshold required for Aβ40 release from perisynaptic astrocytic processes (25–27). This astrocyte-derived Aβ40 could act as the same nucleation trigger described above, shifting the local Aβ42/40 ratio toward the active ∼5% state and producing a delayed-onset activation phase that persists with continued astrocytic Aβ40 delivery into the perisynaptic space.

Because Aβ generation and release operate on slower kinetics than synaptic L-Gln signaling (2), the system may undergo a ligand handover in which early L-Gln–driven activation transitions to Aβ-dependent activation during the functionally sustained phase of LTP, but not its structurally modified phase.

In healthy tissue, the ratio returns to the inactive 10% equilibrium once activity subsides. In MCI, chronic neuroinflammation (30) sustains astrocytic Aβ40 release, maintaining the active ratio and converting a short-lasting physiological signal into a persistent driver of hyperexcitability.

As established above, this active configuration likely enhances INa through vesicular insertion rather than altered gating. As shown for Aβ25–35 (9), the 5% Aβ42/40 state recruits Nav-containing vesicles from intracellular reservoirs such as recycling endosomes, paralleling AMPA receptor insertion during LTP (31, 32). Through microtubule-dependent trafficking (33, 34), the Aβ signal may activate T1R2/T1R3 present in both pre- and postsynaptic regions to trigger presynaptic Nav delivery and postsynaptic AMPA receptor insertion (31, 32, 35), potentially regulating synaptic plasticity such as LTP.

### Part III Mechanism of INa Enhancement and Synaptic Coordination

A central question raised by our findings is how the 5% Aβ42/40 mixture enhances INa amplitude without altering the voltage dependence of activation (Vh). This pattern suggests an increase in the number of functional Nav channels at the plasma membrane, consistent with the behavior of Aβ25–35 (9).

In contrast, Aβ1–40 and Aβ1–42 individually do not produce consistent INa enhancement across Nav1 subtypes; instead, they tend to shift activation toward more negative voltages (36). Thus, the 5% Aβ42/40 mixture behaves functionally more like Aβ25–35, supporting this region as the structural motif relevant to Nav channel modulation. This interpretation aligns with the Aβ40:Aβ42 heterodimer model proposed in Part II, in which the 25–35 region becomes exposed. Such exposure provides a straightforward explanation for why the 5% mixture enhances INa without shifting Vh.

Building on this framework, we propose that dual-ligand activation of T1R2/T1R3 during LTP provides a unified mechanism for activity-dependent increases in synaptic strength. In this model, L-Gln initiates receptor activation during the induction phase, while the Aβ42/40 ratio maintains activation during the later maintenance phase. The resulting continuity of T1R2/T1R3 signaling supports sustained trafficking of Nav-containing vesicles to presynaptic terminals.

Through microtubule-dependent trafficking (33, 34), this Aβ–T1R2/T1R3 axis may coordinate both presynaptic and postsynaptic components of synaptic strengthening. Enhanced microtubule polymerization provides the structural scaffold required to deliver Nav-containing vesicles to presynaptic terminals while simultaneously facilitating the transport of AMPA receptor-containing vesicles to the postsynaptic density (31, 32, 35).

Given that L-Gln/T1R3 signaling is a conserved component of synaptic function from rodent to human (as discussed above), the Aβ–T1R2/T1R3 axis likely adds an additional layer to the synaptic architecture underlying human cognition.

### Part IV. Pathophysiology and Therapeutic Implications

#### A. Chronic Hyperexcitability in MCI

In early AD (MCI), chronic neuroinflammation drives sustained release of monomeric Aβ40 from perisynaptic astrocyte processes. This could persistently shift the local Aβ42/40 ratio toward the active ∼5% state, maintaining chronic activation of the Aβ–T1R2/T1R3 axis. Our findings predict that this sustained activation chronically enhances INa, providing a mechanistic explanation for early neuronal hyperactivity observed before overt neurodegeneration. If this hypothesis holds, it may extend to other neurodegenerative conditions - such as vascular dementia, frontotemporal dementia, and dementia with Lewy bodies - which, despite differing primary pathologies, share common neuroinflammatory mechanisms involving astrocytic and microglial activation (30). This could create a local microenvironment in which astrocytic release of Aβ1-40 into the perisynaptic space is enhanced, driving neuronal hyperexcitability, even though global cerebrospinal fluid biomarker profiles would not reflect a universal systemic rise.

#### B. Concentration-Dependent Transition: Cytoskeletal Collapse and Tau Release

At micromolar concentrations, Aβ peptides disrupt cytoskeletal structure. In our live-cell imaging assays, high Aβ exposure produced loss of filamentous organization in both microtubule- and actin-based structures (36), consistent with cytoskeletal collapse. Such structural disruption would be expected to liberate microtubule-bound tau, generating free tau that can enter the interstitial fluid. This provides a plausible mechanistic basis for the rise of phosphorylated tau as a late-stage biomarker of neurodegeneration (39, 40). This cytoskeletal mechanism naturally aligns with the temporal ordering of Alzheimer’s biomarkers, in which shifts in the Aβ42/40 ratio emerge early while phosphorylated tau rises only after structural collapse has begun.

#### C. A Human-Specific Therapeutic Strategy

Identifying the Aβ–T1R2/T1R3 axis as a uniquely human-sensitive mechanism provides a precise therapeutic window. Rather than broad amyloid clearance, two mechanistically grounded strategies emerge:

1. **Receptor blockade.** Small-molecule antagonists targeting the Aβ binding interface on human T1R2/T1R3 could selectively inhibit Aβ-induced hyperexcitability while preserving L-Gln signaling. Several known sweet-taste inhibitors—including lactisole, dichloroprop, and gymnema extract—suppress INa through T1R3-dependent mechanisms (36). Cinacalcet reduces INa in mouse neurons (41) via T1R3 (36). Anti-T1R3 antibodies inhibit binding of fluorescein-labeled Aβ25–35 (9) and Aβ1–40 (36), suggesting feasibility of therapeutic antibodies or small-molecule screens targeting this interface.
2. **Restoring the physiological Aβ42/40 ratio.** Reducing excess Aβ40 - via Aβ40-specific immunotherapy or Aβ40-selective dialysis—would remove the trigger for chronic pathological T1R2/T1R3 overactivation.

Both strategies converge on suppressing the self-amplifying neuroinflammatory loop (30) that drives early AD progression. Reducing circulating Aβ40 decreases pathological influx of platelet-derived Aβ40 into the brain (42, 43), weakening microglial activation. Blocking Aβ binding to T1R2/T1R3 reduces hyperexcitability, limiting activity-dependent Ca²⁺ elevations in perisynaptic astrocytes that trigger further Aβ40 release (29). By keeping astrocytes below the Ca²⁺ secretion threshold, receptor blockade indirectly dampens astrocyte-driven inflammatory signaling (44, 45). Together, these human-specific strategies offer a rational, mechanism-based path toward stabilizing both molecular signaling and neuronal structure.

These strategies may also have implications beyond early AD symptoms. In late-stage AD, the burden of the disease often shifts more heavily onto caregivers than onto patients themselves. Symptoms such as epilepsy may arise directly from sustained neuronal hyperactivity, while behavioral symptoms such as fear and aggression may reflect a loss of the reference points - of self and surrounding environment - that this hyperactivity disrupts. If so, targeting the Aβ receptor T1R2/T1R3 or restoring the physiological Aβ42/40 ratio could offer meaningful symptomatic relief even when cognitive restoration is no longer feasible, improving quality of life for both patients and their caregivers.

A graphical summary of this conceptual model—from physiological LTP to MCI-associated hyperexcitability—is shown in Figure 5.

**Figure 5.**
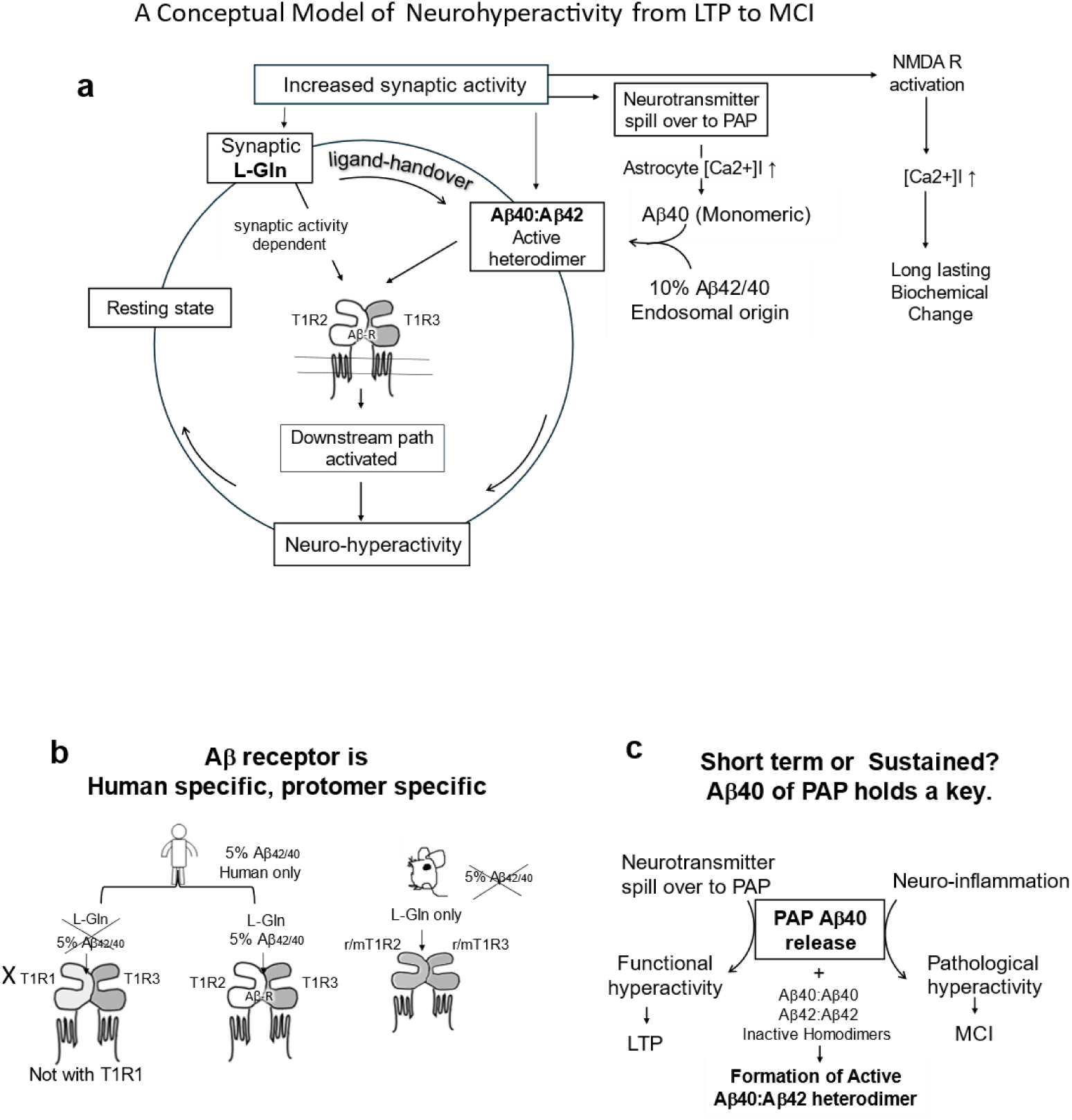
Conceptual Model of T1R2/T1R3 in Synaptic LTP and MCI Progression. (a) Circular diagram illustrating the physiological mechanism of synaptic long-term potentiation (LTP) via T1R2/T1R3. Under heightened synaptic activity, L-glutamine (L-Gln) released from synaptic vesicles activates T1R2/T1R3 and enhances Nav and AMPA receptor (AMPAR) trafficking, while L-glutamate activates ionotropic glutamate receptors, triggering NMDA receptor (NMDAR)-dependent calcium influx leading to biochemical LTP. Concurrently, neurotransmitter spillover to perisynaptic astrocyte processes (PAP) releases Aβ40, which pairs with baseline Aβ42 (10% Aβ42/Aβ40 ratio) to form an active Aβ40:Aβ42 heterodimer (5% Aβ42/Aβ40), which is masked from standard antibody detection. This heterodimer continues to activate T1R2/T1R3 to support synaptic LTP (ligand handover from L-Gln to Aβ), then is replaced with biochemical LTP and returns to basal state to complete the physiological cycle. (b) T1R2/T1R3 is protomer-specific (T1R1/T1R3 does not function as an Aβ receptor; T1R1 knockdown cells behave as wild-type) and species-specific: human T1R2/T1R3 responds to both L-Gln and Aβ, whereas mouse T1R2/T1R3 responds only to L-Gln. (c) The fate of PAP-derived Aβ40 depends on the inflammatory context. Under physiological conditions (left), PAP Aβ40 release maintains functional hyperactivity and LTP. Under neuroinflammatory conditions (right), the same Aβ40 release drives pathological hyperactivity, contributing to mild cognitive impairment (MCI) progression.

## Materials and Methods

### Solutions

The extracellular solution contained (in mM): 126 NaCl, 4 KCl, 1.8 CaCl₂, 1 MgCl₂, 24 HEPES, and 10 d-glucose (pH 7.4, adjusted with NaOH). The intracellular solution contained (in mM): 130 KOH, 130 l-aspartic acid, 5 MgCl₂, 5 EGTA, 4 Tris-ATP, and 10 HEPES (pH 7.2, adjusted with KOH). All chemicals were purchased from Nakalai Tesque (Kyoto, Japan), except for d-glucose, which was obtained from Hayashi Pure Chemical (Osaka, Japan).

The internal patch pipette solution contained K⁺ as the primary cation. Although Cs⁺ is conventionally used to isolate voltage-gated sodium channel (VGSC) currents, K⁺ was retained to preserve the physiological intracellular environment, which is more favorable for intracellular G protein-coupled receptor (GPCR) signaling. This approach may permit outward K⁺ current activation at depolarized potentials, potentially obscuring the VGSC inward current. For this reason, Recordings in which outward current contaminated detection of the VGSC inward current were excluded from analysis.

### Aβ Peptide Preparation

Aβ₁₋₄₀ and Aβ₁₋₄₂ peptide concentrations (5% and 10% Aβ42/40 ratios) were prepared based on values reported by Constantinides et al. (3). Each peptide was dissolved in 10 mM NaOH to prepare a 5 µM stock solution (modified from the Abcam protocol (Abcam. ab120301 Amyloid beta peptide (1–42) (human) protocol., https://www.abcam.cn/ps/products/120/ab120301/documents/Amyloid%20Beta%20Protocol%20-%20LP%2028319%20%28website%29.pdf), yielding a pH of approximately 12.0. Brief sonication was applied to maintain the peptides in monomeric form (47). Because the bath solution volume was 400 µL and 1 µL of peptide solution was applied per experiment, 400× concentrated stock solutions were prepared. Final bath concentrations were: non-demented cohort (NDC): Aβ₁₋₄₂, 254.5 pM; Aβ₁₋₄₀, 2.3 nM; Alzheimer’s disease cohort (ADC): Aβ₁₋₄₂, 123.8 pM; Aβ₁₋₄₀, 2.5 nM. Because the 400× stock solutions yielded a peptide concentration of 2 µM — below the 3 µM threshold at which Nag et al. (48) reported that both peptides remain predominantly monomeric — the stock solutions were considered to be in monomeric form. Peptide solutions were sonicated for 10 s immediately before use and stored at −20°C.

Amyloid β-Protein (1–40) (HCl form, 014-18951) and Amyloid β-Protein (1–42) (HCl form, 011-18961) were obtained from Wako Pure Chemical Industries.

### Aβ Peptide Application

1 µL of the Aβ42/40 peptide mixture (prepared in 10 mM NaOH) was applied to a 400 µL bath solution containing 24 mM HEPES (pH 7.4). The resulting pH change is negligible (ΔpH ≈ 4.2 × 10⁻⁷), as detailed in Saito et al. (9).

### Cell Culture

HEK 293 cells (JCRB Cell Bank), SH-SY5Y cells (ATCC, USA) and NG108-15 (gift from Dr. Furuya, Nagoya University) were maintained in DMEM (Sigma-Aldrich) supplemented with 10% FBS (BioWest, France). NG108-15 cells were additionally maintained in HAT (GIBCO; a liquid mixture of sodium hypoxanthine (5 mM), aminopterin (20 µM), and thymidine (0.8 mM)). For neuronal differentiation, SH-SY5Y cells were treated with 10 µM retinoic acid (Tokyo Chemical Industry, Tokyo, Japan) in DMEM with heat-inactivated FBS for more than 5 days. Although this protocol does not reliably induce long neurites, it reproducibly induces sodium current (INa) expression.

### Stable Cell Line Generation

Nav1.1, Nav1.5, Nav1.6, and Nav1.6/Kv1.2 stable cell lines were generated in HEK 293 or SH-SY5Y host cells. The Nav1.5 expression vector was constructed by subcloning SCN5A (GenBank: BC140813; SourceBioscience, UK) into pcDNA3.1 (Hygro; Invitrogen); BC140813 contains two embryonic exons that were replaced with corresponding exons from human heart cDNA (Zyagen, San Diego, CA), as previously described (9). The following expression vectors were purchased from Genscript and used in this study:

- Nav1.6: Homo sapiens SCN8A, transcript variant 1 (NM_014191.4), pcDNA3.1 Hygromycin
- Nav1.1: Homo sapiens SCN1A, transcript variant 1 (NM_001165963.4), pcDNA3.1 Neomycin
- Kv1.2: pcDNA3.1 Blasticidin

All pcDNA3.1 expression vectors were transfected into HEK 293 and/or SH-SY5Y cells using Lipofectamine 2000 (Thermo Fisher Scientific).

### Transient Expression of hTas1R3 in NG108-15 Cells

hTas1R3 (NM_152228) was cloned into the Vivid Colors™ pcDNA™6.2/EmGFP-Bsd/V5-DEST vector (catalog no. V36620) and transfected into NG108-15 cells using Lipofectamine 2000. Cells were replated onto 8 mm circular cover glasses the following day, and patch-clamp experiments were performed 3–5 days post-transfection. Untransfected NG108-15 cells served as controls.

### T1R1 miRNA Knockdown Vector Construction

The T1R1 miRNA sequence was synthesized and subcloned into the BLOCK-iT Pol II miR RNAi Expression Vector Kit with EmGFP (catalog no. K493600). The shRNA target sequence was selected from the Broad Institute GPP Web Portal; the specific hTAS1R1 (NM_138697) target sequence was CAAGGACACTGTGGCGTTTAA. This sequence was adapted to the BLOCK-iT vector specifications and synthesized as the following SalI/XhoI-flanked segment: gtcgaccagt ggatcctgag gcttgctgaa ggctgta TG CTGTTAAACG CCACAGTGTC CTTGGTTTTG GCCACTGACT GACCAAGGAC AGTGGCGTTT AA caggacac aaggcctgtt actagcactc acatggaaca aatggcccag atctggccgc actcgag

The synthesized segment was cloned into pCDNA6.2-GW-EmGFP-miR at SalI/XhoI sites (Genscript). Successful silencing of T1R1 protein in Nav1.5-expressing HEK293 cells was validated by immunocytochemistry.

### T1R2 CRISPR-Cas9 Knockout

T1R2 knockout HEK 293 cell lines were generated using a CRISPR-Cas9 vector (VectorBuilder). The TAS1R2-targeting guide RNA (5′-GGACCACTCGTTATTCGGGC-3′) was selected using the Broad Institute sgRNA Designer (legacy version) and cloned into the pRP[CRISPR]-mCherry/Puro-hCas9-U6 mammalian expression vector (VectorBuilder, Chicago, IL). This all-in-one vector co-expresses human codon-optimized Cas9, the gRNA (U6 promoter), and mCherry and puromycin resistance markers for selection. Nav1.5 HEK293 cells were transfected with the construct and selected with puromycin to establish the stable KO population. Successful silencing of T1R2 protein in Nav1.5 HEK293 cells was confirmed by immunocytochemistry.

### T1R3 Knockdown Cell Line

The T1R3 knockdown Nav1.5 HEK293 cell line used in this study was established previously, as described in Saito et al. (9).

These genetically modified cell lines (T1R1 KD and T1R2 KO) were subsequently evaluated by immunocytochemistry.

### Immunocytochemistry

Wild-type and knockdown/knockout cells were grown on separate coverslips. During antibody staining, the coverslips were placed side-by-side in separate containers to ensure identical processing conditions. Primary antibodies used were anti-T1R1 (A16788, rat polyclonal, Antibodies.com, Cambridge, UK) and anti-T1R2 (bs-9599R, Bioss Antibodies, Massachusetts, USA).

The staining protocol was adapted from the Abcam immunocytochemistry protocol (https://www.abcam.com/protocols/immunocytochemistry-immunofluorescence-protocol). T1R2 KO cells: Cells were fixed with 4% paraformaldehyde in HEPES-based physiological solution (HBPS) for 10 min at room temperature, washed three times with 400 µL HBPS (5 min each), and blocked with 1% gelatin in HBPS. Primary antibody was applied at 1:4,000 dilution in 1% gelatin HBPS for 1 h at room temperature. After three PBS washes (5 min each), cells were incubated with Cy2-conjugated secondary antibody (Amersham ECL Plex Cy2-conjugated anti-mouse IgG Cy2 (Cytiva, PA42009)) (1% gelatin) for 1 h at room temperature in the dark, then washed with HBPS.

T1R1 KD cells: Because the anti-T1R1 antibody was validated only for western blotting, the standard fixation protocol failed to produce staining. For this reason, paraformaldehyde fixation time was reduced to 2 min and followed by ice-cold ethanol fixation for 5 min to expose the T1R1 epitope by denaturation. A Cy3-conjugated anti-mouse secondary antibody (Amersham ECL Plex Cy3 conjugated anti-mouse antibody (Cytiva, PA43009)) was used.

### Image Acquisition and Analysis

Fluorescence images were captured using an IDS UI-3880CP-M-GL R2 camera mounted on the microscope and acquired with uEye Cockpit software (IDS Imaging Development Systems GmbH). Images were analyzed using ImageJ 1.554f (49) and annotated in Microsoft PowerPoint. When direct fluorescence intensity comparisons were made between control and experimental groups, all images were processed identically.

### Microscopy and Patch Pipette Manipulation

Cells were grown on uncoated round cover glasses (8 or 12 mm diameter) placed in a recording chamber filled with 400 µL of extracellular solution. Cells were visualized using a phase contrast inverted microscope (Olympus IMT-2) with a ×40 objective and ×10 eyepiece. Patch pipettes were pulled from borosilicate glass capillaries (CG150F-10; Harvard Apparatus) using a PC-10 puller (Narishige) and had resistances of 3–5 MΩ. A water hydraulic micromanipulator (WR-91; Narishige) was used to position the pipette and achieve gigaseal formation under visual observation.

### Whole-Cell Voltage-Clamp Electrophysiology

Patch-clamp recordings were performed using an Axopatch 200A amplifier (Axon Instruments, Union City, CA). Signals were digitized with a Digidata 1332A interface and acquired using Clampex (pClamp 9.0; Axon Instruments). Currents were filtered at 5 kHz and digitized at 10 kHz. Cells were held at −120 mV, except for Nav1.6-expressing cells, which were held at −80 mV. Sodium currents were elicited by voltage steps from −80 mV to +50 mV in 10-mV increments (I–V protocol). NG108-15 recordings were obtained from large round cells to facilitate giga-seal formation.

Following the I–V protocol, an I–T protocol was performed. In this protocol, the membrane was held at −80 mV, stepped to −120 mV for 1 s, and then depolarized to −20 mV to activate INa. This sequence was repeated every 10 s. Once the INa amplitude reached a stable baseline, 1 µL of peptide (Aβ42/40 mixture) solution was applied to the bath. Peptide effects were assessed after a minimum of 4 min of repeated I–T cycles. After peptide incubation, the I–V protocol was repeated to evaluate changes in INa enhancement.

For Nav1.6-expressing cells, a holding potential of −80 mV was used because deeper hyperpolarization reduced functional INa availability. The −80 mV holding potential prevented cumulative inhibition to evaluate the effect of peptide.

During the I–T protocol, if rundown of INa was observed before peptide application, the control I–V amplitude was adjusted by the same amount to ensure accurate comparison with the post-peptide I–V curve. Series resistance compensation was not employed. Leak subtraction was applied during analysis to remove the linear leak component.

To evaluate peptide effects, the negative peak amplitude of the control INa was normalized to −100 (or 1), and post-peptide values were expressed relative to the normalized control value.

Peptide-induced enhancement of INa was quantified as Percent Enhancement, calculated for each cell as:

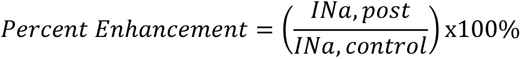

Each I–V curve was converted to a conductance–voltage (G–V) relationship, and half-activation voltages (Vh) were obtained by fitting to the Boltzmann equation:

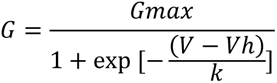

Because voltage steps were applied in 10-mV increments, the slope factor k could not be reliably determined from individual cells. Therefore, only Vh was extracted for each cell, and k was estimated only from averaged G–V curves.

The difference in half-activation voltage (Vh diff) was calculated as:

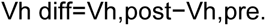

Averaged I–V plots are presented as means ± SE. Data were analyzed using Clampfit 9.0.1.07, and plots were generated in Microsoft Excel. Boltzmann and Hill equation fitting was performed using CurveExpert Professional (Hyams Development).

### I-V plot construction

When generating the averaged I–V curves, voltage offset variations among cells in the control recordings were corrected for each cell to ensure control I–V consistency across all tested cells, and the corresponding correction was applied to the peptide condition to maintain consistency between control and peptide-treated traces. Because the K Asp internal solution—chosen to model physiological GPCR signaling conditions—permitted outward IK activation at depolarized potentials, those outward data points—although recorded—were excluded from the averaged I–V curves when IK became large enough to distort the INa profile. Only the inward portion was used for G–V curve Boltzmann equation fitting, as peak INa, Vh, and slope factor are determined solely from the inward activation range. Therefore, the n values reflect only the number of cells contributing data within the INa activation-to-peak range.

### Statistical Analysis

The Mann–Whitney U test was used to assess the significance of peptide-induced changes in INa amplitude. Statistical analyses were performed using the SciPy library in Python (50). A p-value < 0.05 was considered statistically significant. Data are presented as mean ± SEM.

## Acknowledgments

During the preparation of this manuscript, the authors used Claude and MS Copilot to improve the language and flow of the text. After using these services, the authors reviewed and edited the content as needed and take full responsibility for the final version of the manuscript.

## Author Contributions

M.L.S. designed research, performed research, analyzed data, and wrote the paper. M.R.S. performed visualization, reviewed and edited the manuscript, and administered the project.

